# A Single-Cell Atlas of Transcriptional and Immunoglobulin Repertoire Evolution in Early B Cell Development

**DOI:** 10.1101/2025.10.08.681095

**Authors:** Charlotte DiBiase, Jonathan Hurtado, Bryan Briney

## Abstract

The development of human B cells in the bone marrow can be separated into functionally and transcriptionally distinct subsets, which while canonically described have yet to be profiled deeply and individually and with NGS and bioinformatic techniques. In this study, single-cell RNA sequencing was performed on 65,110 B cells from six healthy donors to observe the foundation of the diversity observed in mature immune repertoires. Following the established narrative of development, committed B cells undergo heavy and light chain recombination, which together form the functional BCR. However, a granular per-subset phenotypic analysis reveals proliferative bursts following each recombination event, an aspect of BCR recombination previously undescribed in the pro-B phase of development. Heavy and light chain pairing becomes more similar to that of mature, circulating B cells with progress through lymphopoiesis, deleting features associated with autoreactivity. Prominent among these repertoire alterations is a substantial shortening of heavy chain CDR3s, and changes in V, D, and J gene usage. This study provides a detailed snapshot of the transcriptomic and repertoire genetics landscape of developing B cells, providing a deeper understanding of the repertoire-shaping influence of early selection processes, and introducing new avenues for repertoire development research.

## INTRODUCTION

An individual’s B cell repertoire originates in the bone marrow as hematopoietic stem cells (HSCs) which differentiate, recombine and mature throughout a lifetime into some combination of a possible 10^18^ unique antibodies (Briney et al. 2019). Their immense generated diversity can be attributed to various elements of development, maturation, and exposure. When cells exit the bone marrow as transitional B cells, they have fully functional B cell receptors (BCRs), which are able to recognize antigens via their antigen binding site (Loder et al. 1999; Nemazee 2017). B cell receptors are composed of heavy (HC) and light (LC) chains, which are generated independently (Pieper, Grimbacher, and Eibel 2013). Heavy chain regions include variable (V), diverse (D), and joining (J) segments coded by ∼51 IGHV, 27 IGHD, and 6 IGHJ gene segments, respectively, which in different combinations contribute to overall repertoire diversity (Matsuda et al. 1993; Corbett et al. 1997; Ichihara, Matsuoka, and Kurosawa 1988). Light chains lack D gene segments and are classified as kappa light chains (KLC, 40 IGKV, 5 IGKJ) or lambda/λ light chains (LLC, 30 IGLV, 7 IGLJ) (Barbié and Lefranc 1998; Pallarès et al. 1998).

The formation of the BCR spans multiple stages of early B cell development. Once cells have committed to the B cell lineage from HSCs in the the pre pro-B phase (marked by downregulation of FLT3 and upregulation of IL7-R and EBF1), they enter the pro-B phase (Kikushige et al. 2007; Kaiser et al. 2023; Schwickert et al. 2014). There, the heavy chain rearranges itself via double strand DNA breaks mediated by RAG1/2, first recombining the D and J segments, and if successful, the V segment to the recombined D-J (Chi, Li, and Qiu 2020; Bassing, Swat, and Alt 2002; Tonegawa 1983). Ligation of these regions of the BCR HC is accompanied by exonuclease-mediated trimming and N-addition, the non-templated addition of nucleotides, at the junction of D-J and V-D segments. N-addition is facilitated by DNTT/TdT and is responsible for generating the majority of pre-immune BCR repertoire diversity (Bassing, Swat, and Alt 2002; Tonegawa 1983; Bendall et al. 2014). Proper folding of the newly recombined μHC is tested by pairing with the surrogate light chain (composed of VpreB and λ5 segments) to form the pre-BCR (Khass et al. 2016; Winkler and Mårtensson 2018; Melchers 2015). Successful formation of a functional pre-BCR is a major checkpoint for B cell development, triggering expression of CD79A/B and CD19, leading to large scale proliferation marked by CD20/MS4A1, MK167, PTMA, and PCNA expression in the next developmental phase, large pre-B (Fuentes-Pananá et al. 2006; Morgan and Tergaonkar 2022; Lee et al. 2021). The final step of BCR assembly is LC rearrangement, which happens in small pre-B cells. Recombination is attempted first at the KLC locus, and LLC rearrangement occurs only if the rearranged KLC fails to pair with the μHC, is misfolded, or autoreactive (Collins and Watson 2018). If both KLC and LLC rearrangement fail, the cell undergoes apoptosis mediated by BCL7A and GADD45A (Okoreeh et al. 2022; Nemazee 2017). Once HC and LC are successfully paired and form a functional BCR in the intermediate B phase, they exit the bone marrow for further maturation and eventual antigen exposure (***Figure 1***).

**Figure 1.**
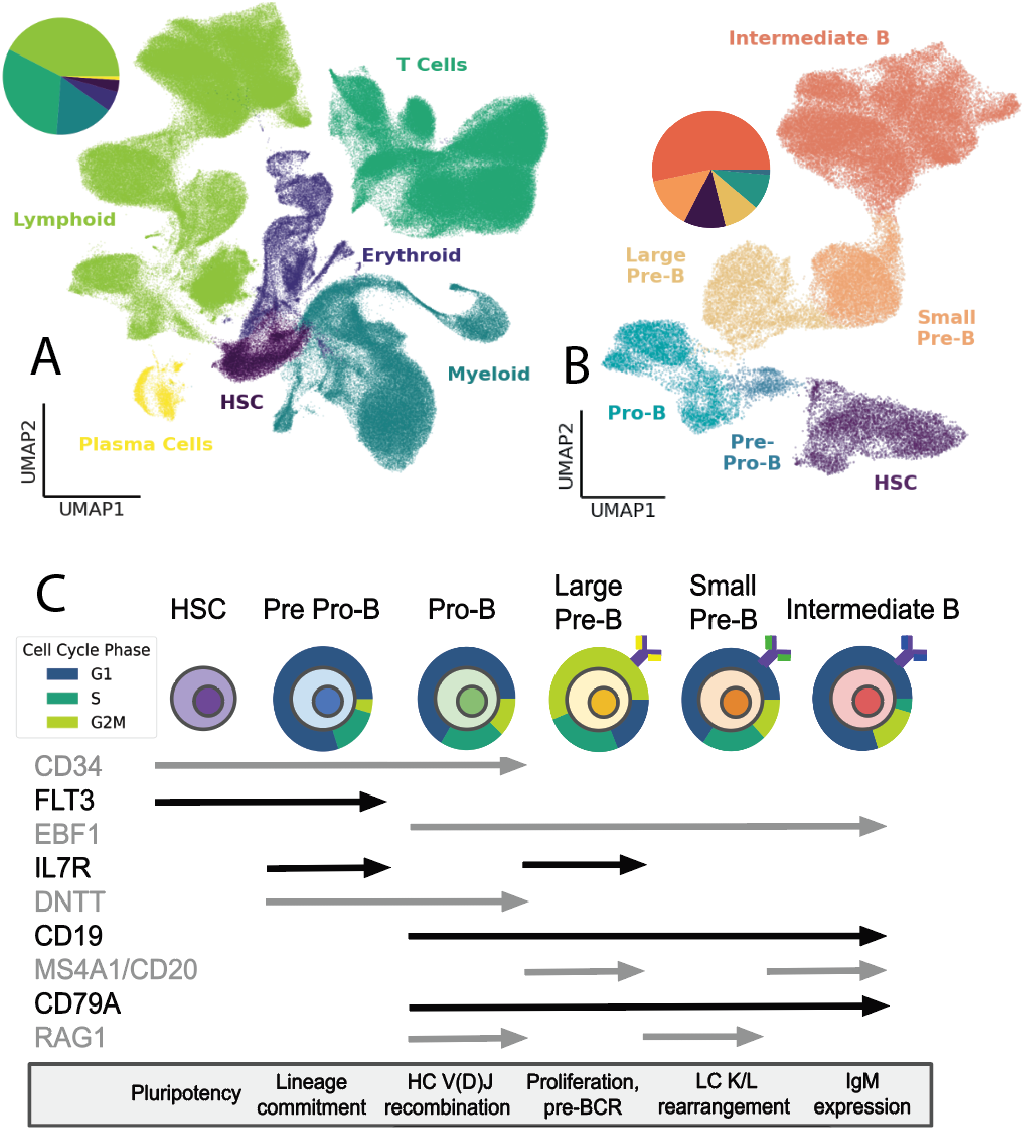
Stages of early B cell development. **(a)** Full lineage UMAP of 388941 harvested immune cells containing 125,153 myeloid cells (32.5%), 111,500 lymphoid cells (28.8%), 100,327 T cells (25.9%) 45,766 erythroid cells (11.8%) 2,381 plasma cells (0.6%) and 1,276 HSCs (0.4%) **(b)** B cell UMAP of 65,110 B cells clustered into 34,704 intermediate B cells (53.3%), 9,241 small pre-B cells (14.2%), 7,458 HSCs (11.5%), 6,366 large pre-B cells (9.8%), 6,348 pro B cells (9.7%), and 993 pre-pro B cells (1.5%). **(c)** Expression of marker genes throughout B cell development alongside major developmental events at each stage. Expression continues through the length of each arrow. HC is shown in purple, SLC in yellow, rearranging LC in green, and final LC in blue. Proportions of cells in each cell cycle phase are shown around each cell type (pre pro-B: 79.5% G1, 14.6% S, 5.9% G2/M, pro-B: 66.3% G1, 21.6% S, 12.1% G2/M, large pre-B: 18.8% G1, 25.0% S, 56.2% G2/M, small pre-B: 65.5% G1, 21.4% S, 13.1% G2/M, intermediate B: 80.0% G1, 4.4% S, 15.6% G2/M).

The BCR’s third complementarity determining region (CDR3) encompasses portions of the V and J regions as well as the entire D region in HCs, and is the primary determinant of an antibody’s antigen specificity (Akbar et al. 2021). The length of the CDR3, particularly of the heavy chain (CDRH3), is often considered in analysis of BCR development because longer sequences have an increased propensity for autoreactivity due to their increased flexibility and tendency to encode multiple hydrophobic amino acid residues (Xiao et al. 2024; Guloglu and Deane 2023). Because longer CDRH3s are more likely to be autoreactive, they are preferentially weeded out at various points in development, most often at the pre-BCR checkpoint between large pre-B and intermediate B phases (Khass et al. 2016). Thus, it has been long understood that the CDRH3 length distribution amongst developing B cells shortens during development, however the field lacks clear examples of this in robust human datasets (Mejias-Gomez et al. 2023). Metrics of human immune repertoire diversity include traits determined early in development such as V_H_/J_H_, V_H_/D_H_ or V_H_/V_L_ clonotypes, and CDRH3 length (Guo et al. 2025; Briney et al. 2019). Here, we aim to understand what the transcriptomic landscape of developing B cells reveals about the origin of immune repertoire diversity through examination of BCR clonotype dynamics, CDRH3 length, and gene regulatory networks.

## METHODS

### Sample preparation and sequencing

Bone marrow samples from four healthy adult donors were obtained through our commercial clinical partner (Hemacare, now Charles River). Lymphocytes were isolated by density gradient centrifugation using Lymphoprep (StemCell Technologies). To ensure representation of early progenitors and developing B cell populations, donor samples were individually enriched for B cells and hematopoietic stem cells by EasySep Human B Cell Enrichment and Human CD34^+^ Selection Kits (StemCell Technologies).

Single-cell RNA sequencing (scRNAseq) was performed on over 3.8×10^5^ cells from bone marrow and enriched samples with the 10x Genomics 5’ Single Cell Immune Profiling Kit (v2). Briefly, donor samples were individually labelled with Totalseq Anti-Human Hashtags (Biolegend) and pooled prior to overloading the 10x Genomic reaction, targeting 4-6×10^4^ cells per run (Stoeckius et al. 2018). Single-cell emulsion generation, barcoding, V(D)J enrichment, and sequencing library construction were carried out according to theChromium Next GEM Single Cell 5’ Reagent Kits v2 (Dual Index) protocol (CG000330 Rev A). Final libraries were sequenced on a NovaSeq 6000 using a 26-10-10-90 read configuration, with minimum target sequencing depths of 5,000 reads per cell for Cell Surface Protein libraries, 10,000 reads per cell for V(D)J libraries, and 30,000 reads per cell for Gene expression libraries.

### Data processing

FASTQ files were generated from raw base call files using CellRanger, and downstream analyses were performed with scab (Hurtado et al. 2022). Matrices of VDJ contigs and counts were assembled by alignment to VDJ and transcriptome reference genomes, outputting full-length per cell VDJ sequences as FASTA files and expression profiles (cells x genes) as an .h5 file, differentiating individual cells by barcode. Antibody sequences were annotated using abstar (Briney and Burton 2018) and integrated with gene expression and cellhash data using scab. Donors were demultiplexed by cellhash using scab.

Because samples had different read depths, cell hash thresholds were incorrectly high for some of the samples with fewer reads per cell. Reads were thus normalized to ensure equal representation from each donor. To further refine the expression matrix, cells with fewer than 200 and greater than 6000 genes were removed. Genes which appeared in fewer than 50 cells were also removed. Normalization, log plus one transformation, and scaling were performed using scab and scanpy. Scab’s dimensionality reduction tool performed principle component analysis, neighborhood graphing, and cell clustering according to the Leiden algorithm and utilized UMAP for embedding (Traag, Waltman, and van Eck 2019; Becht et al. 2018; McInnes, Healy, and Melville 2018). Doublets were identified using Scrublet (Wolock, Lopez, and Klein 2019) and all doublets, including entire clusters containing primarily doublets, were removed.

Cells were then re-clustered without doublets and identified as lymphoid, myeloid, erythroid, plasma cells, T cells, or B cells according to gene expression profiles. B cells were then separated out, clustered again for granularity, and subset into developmental phases also according to gene expression profiles.

### Bioinformatic analyses

To explore transcription of canonically relevant developmental genes, umap plots were generated using scab, and seaborn’s boxenplot function was used to further visualize expression level distributions between subsets (Waskom 2021). Sankey diagrams of heavy chain VDJ segment frequencies were visualized using plotly. Other visualizations were performed in Python using seaborn and matplotlib (Hunter 2007; Waskom 2021).

Paired heavy and light variable region amino acid sequences were analyzed using ImmunoMatch to determine pairing score distribution differences between subsets (Guo et al. 2025). A one-versus-rest support-vector-machine (SVM) was used to classify subsets from V_H_/D_H_ and V_H_/V_L_ clonotypes. Classifier training, using a combination of V_H_, V_L_, and J_H_ genes along with CDRH3 length, was performed in Python using Scikit-learn (Pedregosa et al. 2011).

Differential gene expression analysis was performed to identify genes most differentially regulated between subsets using the rank_genes_groups function in scanpy (Wolf, Angerer, and Theis 2018). The top 50 most differentially expressed genes in each subset were used for ontology analysis using g:Profiler (Kolberg et al. 2023). Enriched terms were selected based on adjusted p-value. DEG analysis before and after pre-BCR formation was performed separately in R using DESeq2 (Love, Huber, and Anders 2014), and ontology analysis of the 28 genes isolated was performed with enrichR (Xie et al. 2021). Pro-B differentially expressed genes were also analyzed with a boruta random feature selection and SHAP analysis to determine genes contributing to subset classification (Kursa and Rudnicki 2010; Lundberg and Lee 2017). Functional profiling of proteins from DEGs was visualized with STRING db (Szklarczyk et al. 2023).

## RESULTS

### Immune cell landscape profiling and B cell developmental stage identification

Transcriptional information from 453,615 scRNA-sequenced bone marrow derived cells was preprocessed, filtered, and normalized (see ***Methods*** for preprocessing details), resulting in 388,941 immune cell profiles from four donors. Transcriptionally distinct clusters were identified as separate immune lineages using known marker genes: CD34 (hematopoietic stem cell; HSC), CD3D (T cells), CD14 (myeloid lineage), HBB/ANK1/EPOR (erythroid lineage), MS4A1/PAX5 (lymphoid lineage), and CD27 (plasma cells). B cells comprised 65,110 of the total cell population. Within B cells, 19 distinct clusters were identified which were subset into six known developmental stages: HSC, pre pro-B, pro-B, large pre-B, small pre-B, and intermediate B. Alongside surface and intracellular markers associated with these stages of development, markers of mechanistic processes characteristic of certain developmental phases, such as V(D)J combination (RAG1/2) in the pro-B phase, were used to validate subset assignments.

### Developing B cells cycle between recombination and proliferation

We observed a prominent phenotypic signature of early B cell development to be alternating cycles of recombination and proliferation. Proliferative bursts, as identified by increased expression of activation and proliferation markers like PTMA, PCNA, and MKI67 (***Figure 2A***), immediately follow each heavy chain recombination event as identified by increased expression of recombination markers like RAG1/2, DNTT and ADA (***Figure 2C***). Upregulation of apoptosis markers was observed alongside recombination markers (***Figure 2B***), presumably an indicator of unproductive recombination. Post-recombination proliferation is further supported by cell cycle analysis showing phase transitions consistent with replication concurrent with upregulation of the aforementioned activation and proliferation markers (***Figure 2E***). While it has previously been shown that pre-BCR signaling triggers expansion in pre-B cells (Lee et al. 2021), we also observed proliferation following D-J recombination (***Figure 2E***). This is particularly interesting because, unlike pre-B cell expansion initiated by a functional pre-BCR, there is no clear mechanism for direct signaling by the products of a successful D-J recombination event. DEG analysis comparing proliferating vs recombining pro-B cells suggest JCHAIN may be directly or indirectly involved, as it is upregulated as D-J recombination is completed and directly prior to the post-recombination proliferative burst.

**Figure 2.**
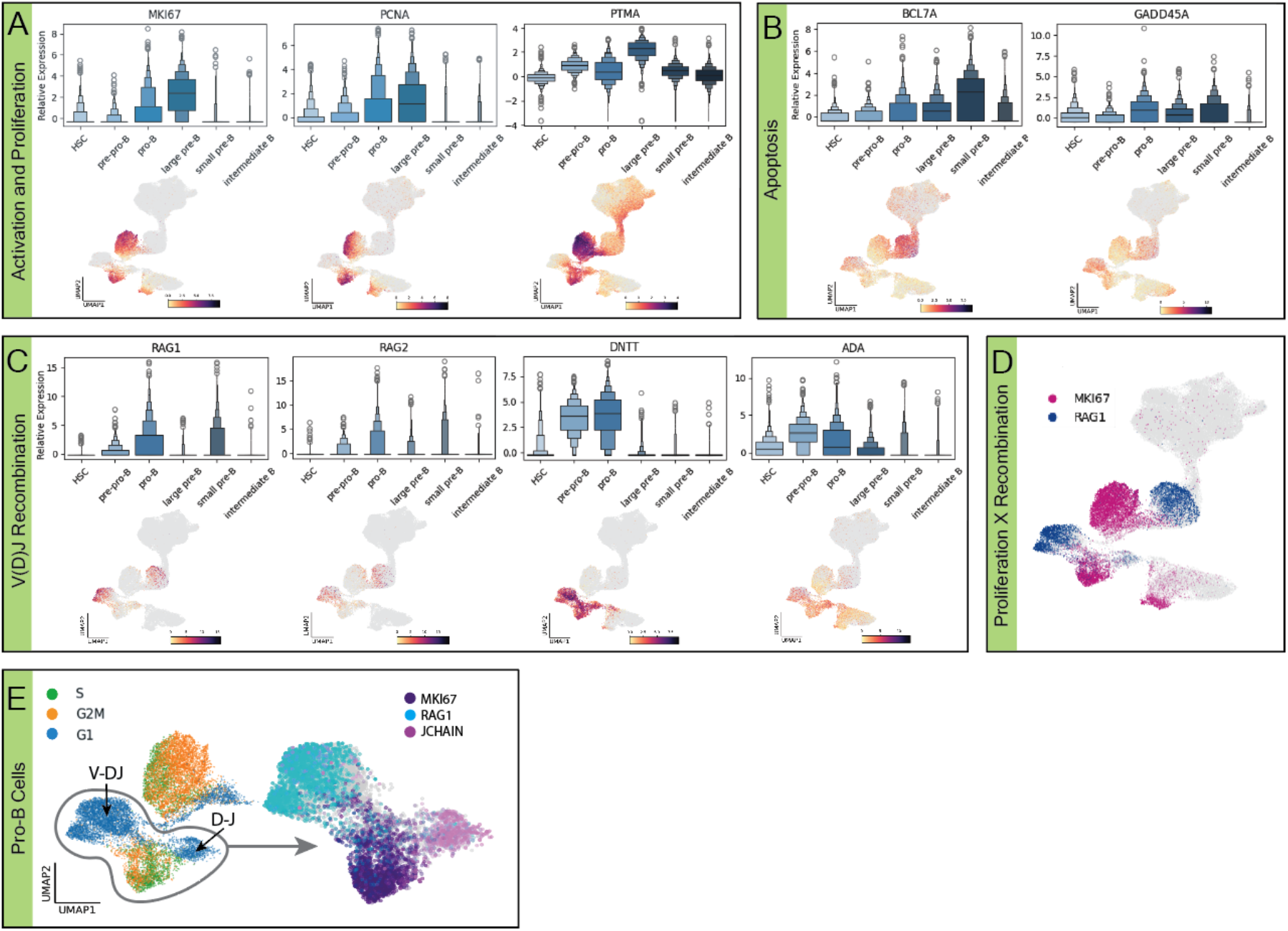
Developing B cells alternate between recombination and proliferation. Expression boxenplots by subset and full B cell lineage UMAPs grouped by biological processes **(a)** activation and proliferation **(b)** apoptosis, and **(c)** V(D)J recombination, relative expression shown by point shade. **(d)** Coexpression UMAP of proliferation (MK167) and recombination (RAG1), relative expression is indicated by color intensity. **(e)** Cell cycle progression and coexpression of proliferation/recombination markers in the pro-B subset.

### Antibody repertoire dynamics across developmental subsets

Heavy chain rearrangement in the pro-B phase is represented by the initiation of expression of V and J genes in isolated heavy chains. The likelihood of pairing between each V or J gene did not vary significantly across subsets, suggesting the recombination process itself, rather than subset-specific selection processes, are the cause of skewed V and J gene use (***Figure 3A–B***). The distribution of CDR3 lengths changes markedly across early B cell development, as heavy and light chains come together form the BCR and central tolerance mechanisms eliminate autoreactive clones. Unique CDR3 sequences from 6178 Large pre-B, 8152 small pre-B, and 27096 heavy chains were investigated for variation in length distribution. Mean amino acid length for large pre-B, small pre-B, and intermediate B were 17.891, 17.615, and 15.422, respectively. Interquartile ranges were 6, 7, and 5 amino acids (***Figure 3C***). Frequency of CDR3 sequences including shared clonotypes was similar to distribution of unique CDR3 sequences.

**Figure 3.**
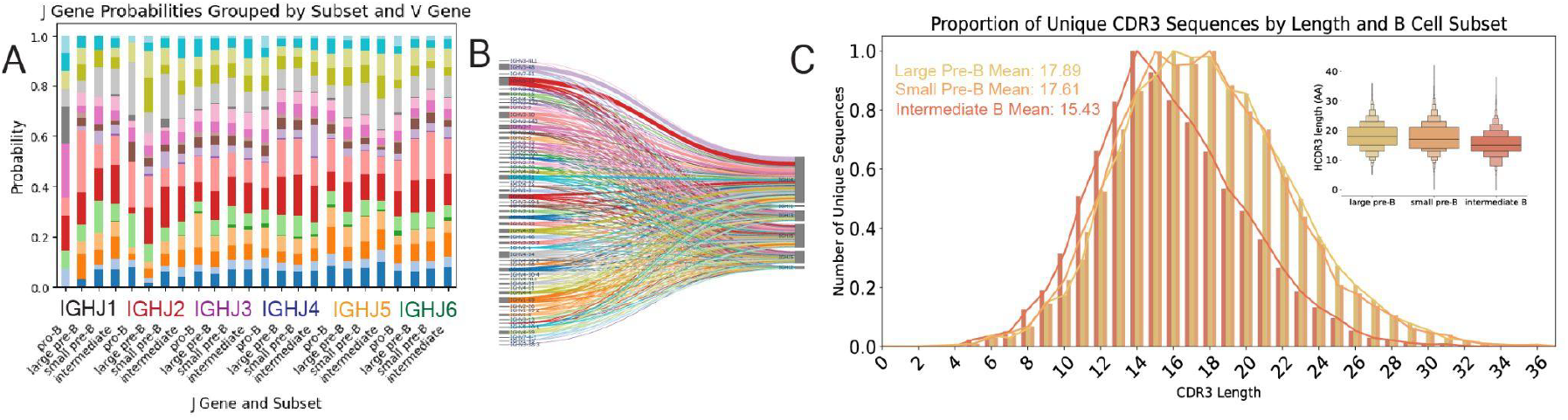
(a) Probability of the combination of J and V genes in individual cells in each developmental stage after V(D)J recombination occurs. Colors correspond to V genes from the Sankey diagram. (b) Sankey diagram of V and J combinations across developmental stages. V genes are differentiated by color, each line representing a single cell. (c) CDRH3 sequence lengths in small pre-B, large pre-B, and intermediate B. Inlay shows boxenplot of length distributions. Mean CDR3 lengths from each subset are large pre-B: 17.89, small pre-B: 17.61, and intermediate B: 15.43.

### ML for V(D)J sequence analysis between subsets

We employed several machine learning (ML) techniques to identify genetic features that distinguish the expressed antibody repertoires of each developmental subset. Pairing score distributions were calculated using Immunomatch (Guo et al. 2025) on V_H_/V_L_ pairs from 947 large pre-B cells, 2,229 small pre-B cells, and 11,136 intermediate B cells, showing the similarity between amino acid sequences from developing B cells to the mature B cells the model was trained on. All three subsets have bimodal distributions centered around 0 (low probability of pairing) and approximately 0.7. Distributions shift towards higher pairing scores as cells progress through lymphopoiesis, with large pre-B cells having the highest peak at 0, small pre-B having relatively equal peaks, and intermediate B cells being most skewed towards high scores, with an additional peak at approximately 0.9 (***Figure 4A***). SVM modeling was used to predict the developmental subset from V(D)J region amino acid sequences of heavy and light chains. Using the D and V regions of heavy chains, pairs from large pre-B, small pre-B pairs, and intermediate B had AUCs of 0.55, 0.51, and 0.57, respectively (***Figure 4B***). Using the V region sequences of heavy and light chains, pairs from large pre-B, small pre-B pairs, and intermediate B had AUCs of 0.64, 0.61, and 0.6, respectively (***Figure 4C***).

**Figure 4.**
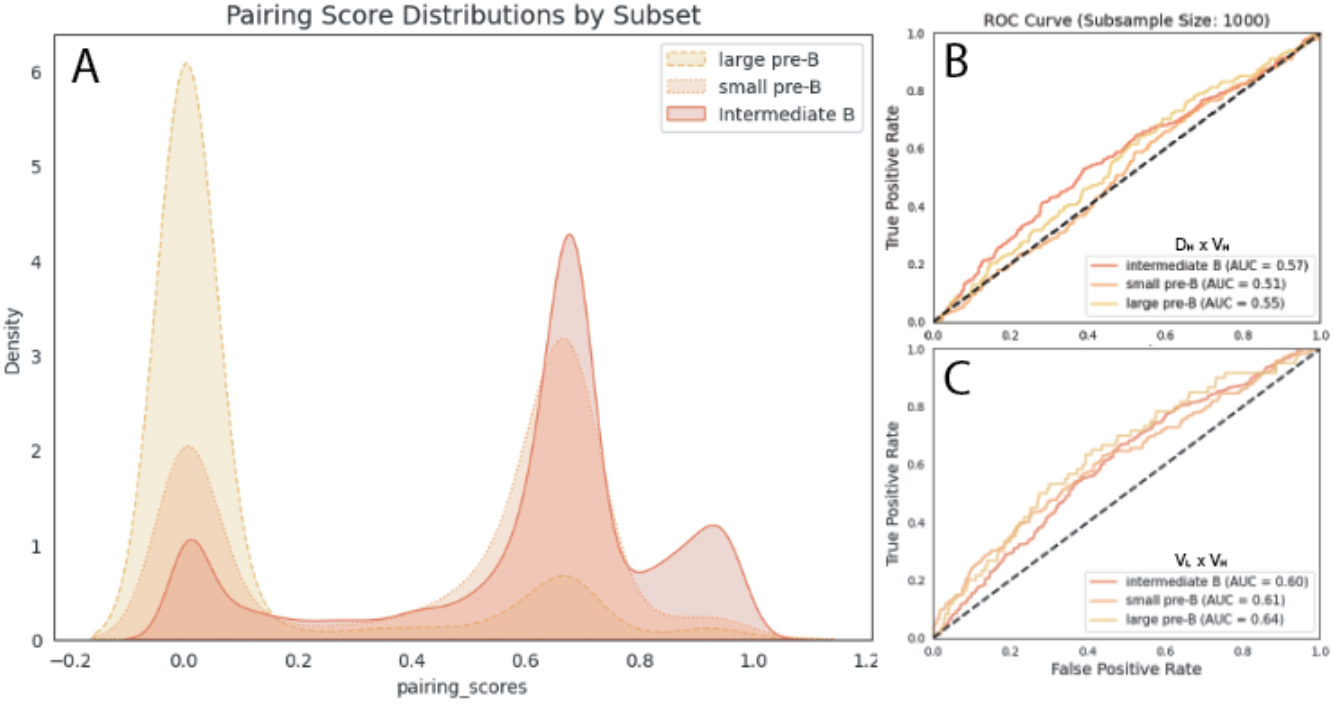
(a) Distribution of pairing scores of V_H_/V_L_ pairs calculated using ImmunoMatch (b) AUC-ROC curve from SVM model trained on D_H_/V_H_ and (c) V_H_/V_L_ pairs from small pre-B, large pre-B and intermediate B subsets.

### Ontology analysis

Gene ontology analysis was performed on the top 50 most differentially expressed genes from clusters associated with each subset of development (***Figure 5***). Enriched ontology terms throughout developmental stages reflect the stages and checkpoints of B cell initialization. In the HSC stage, enriched terms are pertaining to metabolic processing (transketolase activity, organonitrogen compound metabolic process), and ribosomal activity (rRNA binding, structural constituent of ribosome, cytosolic small ribosomal subunit). Pre-pro B cell expression was enriched in processes surrounding immune pathway differentiation (leukocyte proliferation, regulation of mature B cell apoptotic process, immune system process, regulation of response to stimulus, intracellular signaling cassette) and signaling pathways (protein binding, regulation of cell motility, extracellular exosome, cytoplasmic vesicle lumen). In pro B cells, ontology terms related to pre-BCR expression (cellular biosynthetic process, immune system process, intracellular signal transduction, response to stimulus).

**Figure 5.**
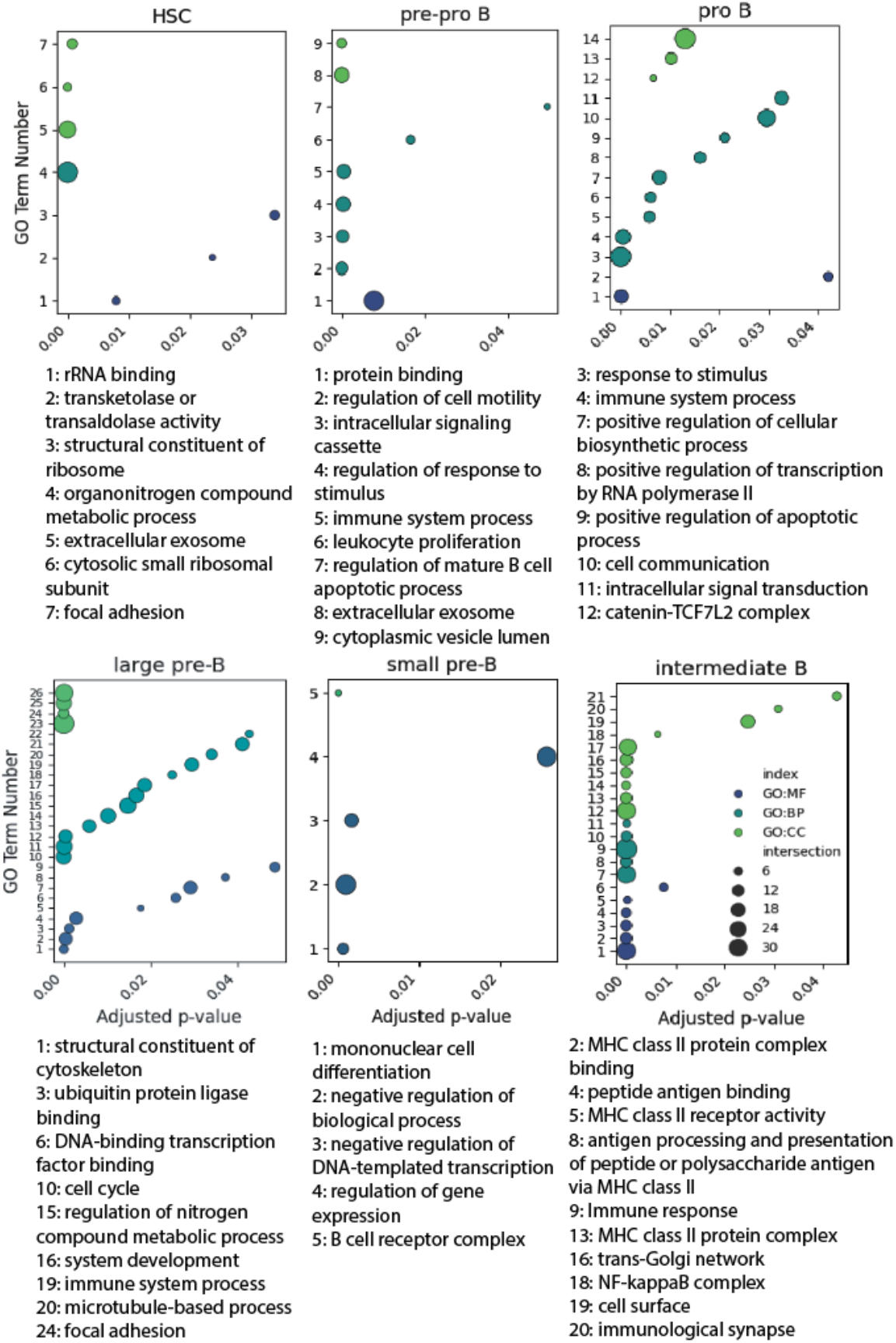
Enriched ontology terms in developing B cell subsets. Molecular function, biological process, and cellular component related terms are shown in green, teal, and blue, respectively. Relative enrichment is measured by -log10(adjusted P) value of each term relative to all others considering the 50 most differentially expressed genes in that subset. Full table of enriched terms in supplementary figures.

Large pre-B cell expression was enriched in clonal expansion (cell cycle, regulation of metabolic process, structural constituent of cytoskeleton, and various protein and transcription factor binding). In small pre-B cells, enriched ontology terms surrounded light chain rearrangement (negative regulation of biological process and DNA-templated transcription) and the newly fully functional BCR (B cell receptor complex). Lastly, in intermediate B cells, many enriched terms were pertaining to IgM expression (MHC class II receptor activity and protein complex, NF-kappaB complex, immune response, antigen processing and presentation).

### Differential expression around pre-BCR formation

27 genes were found to be differentially expressed pre- and post-BCR formation (***Figure 6A***). Proteins associated with these genes clustered into functional profiles related to positive regulation of exit from mitosis, bladder cancer, DNA replication dependent chromatin assembly, lambda 5 deficiency/DNA recombinase complex. Most enriched ontology terms across all GO pertained to regulation of cell cycle (G1/S transition of mitotic cell cycle, DNA integrity checkpoint signaling) and protein synthesis (lumenal side of endoplasmic reticulum, intracellular membrane bound organelle) (***Figure 6B***). Analysis of differentially expressed genes by heatmap (***Figure 6C***) or volcano plot (***Figure 6D***), showed most were expressed more highly post-recombination, suggesting successful recombination initiates increased transcription of genes across multiple biological pathways.

**Figure 6.**
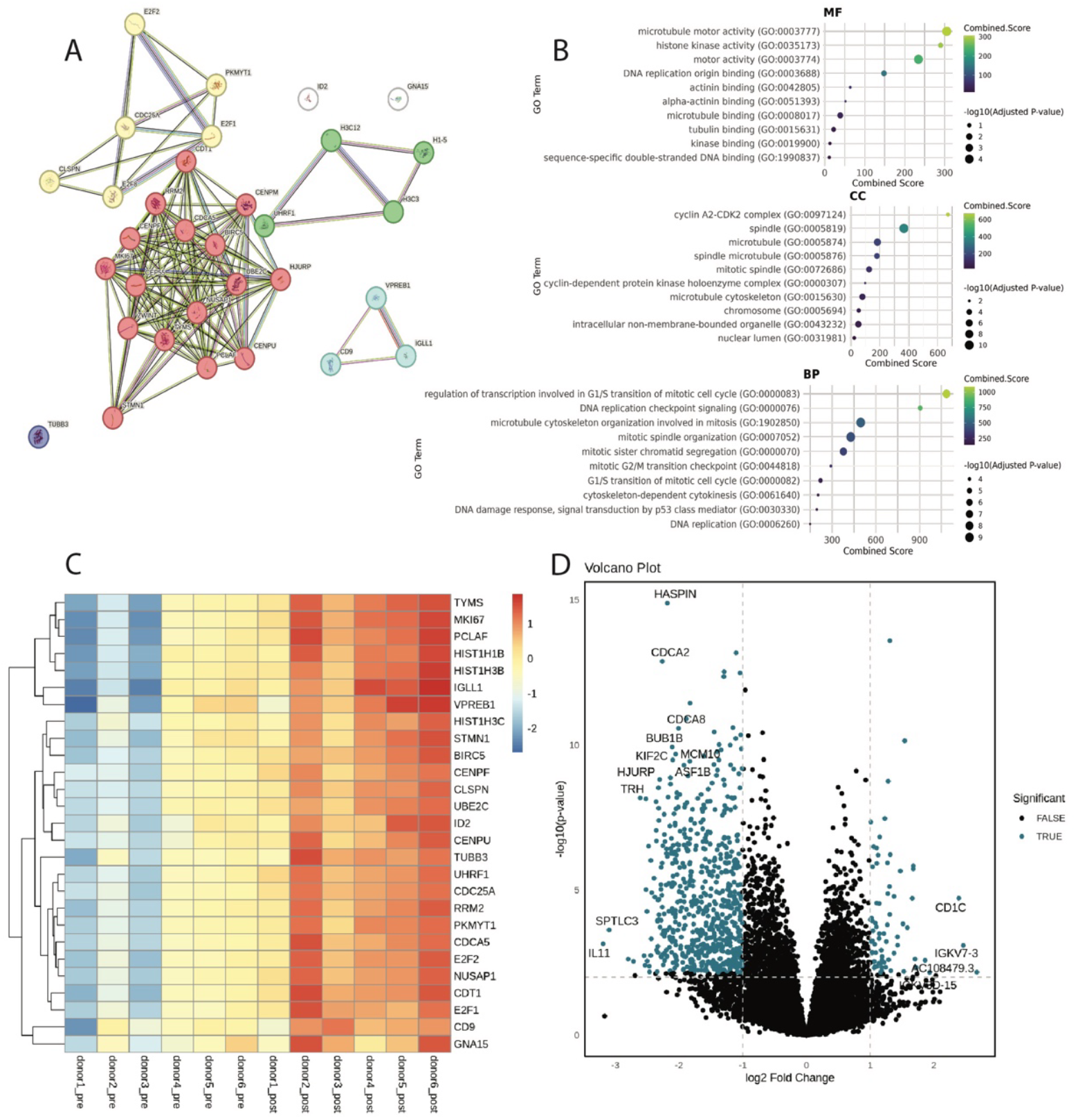
Differentially expressed genes before and after the formation of the pre-BCR. (a) STRING functional protein association network, red: positive regulation of exit from mitosis, yellow: bladder cancer, green: DNA replication-dependent chromatin assembly, blue: DNA recombinase complex, purple: TUBB3 (b) ontology analysis of molecular function (MF), cellular component (CC) biological process (BP) and related enriched terms (c) Heatmap of differentially expressed genes before (pre) and after (post) pre-BCR formation across donors. (d) Volcano plot of genes expressed before and after pre-BCR formation.

## DISCUSSION

As developing B cells mature, the genes and features present are continuously assembled and recombined. This filtering and shuffling of the antibody repertoire impacts autoreactivity and functionality. Here, the gene expression dynamics of repertoire refinement are categorized into previously defined developmental subsets, detangling the relationships between stages of B cell lymphopoiesis and the construction of the naive antibody repertoire.

### Shortening of CDRH3 implies regulation of autoreactivity and optimization of antigen binding

During V(D)J recombination, the CDRH3 sequence is determined by the combination of V, D, and J genes from which it is made up. Because excessively long CDR3 sequences threaten autoreactivity of the BCR, the length of sequences throughout development reflects how heavy chain rearrangement and downstream mechanisms of central tolerance mitigate self-reactivity (Wardemann et al. 2003). Reduction of CDRH3 length by 2.2 amino acids on average between small pre-B cells and intermediate B cells shows how the process of light chain recombination may favor shorter CDRH3 sequences in the formation of the BCR, likely to increase structural viability and reduce risk of autoreactivity in mature B cells (Figure 3c). A study of 9684 paired mature B cell sequences found average CDRH3 length to be 14.54 amino acids, suggesting that downstream self-regulation continues to favor shorter sequences (from the 15.43 average length calculated here) (Mejias-Gomez et al. 2023). CDRH3 length can also play a role in immunogen engagement and thus shortening of the sequence throughout development could also reflect a tendency of antibody CDRH3 regions to more closely resemble their respective potential antigens (Swanson et al. 2023). This observed reduction of length is one of the first concrete examples of the process of B cell maturation favoring shorter CDRH3 sequences and provides the precise developmental timing of the shortening process, allowing for more in depth future investigation.

### Alternation of recombination and proliferation demonstrates movement through viability checkpoints before replication events

Expression patterns of proliferation, recombination, and apoptosis genes highlight the largely stepwise process of B cell development. While alternation of proliferation and recombination have been previously suggested, these data strongly imply a largely mutually exclusive relationship between these two processes (Hamel et al. 2014), Figure 2d). Previous research has focused on individual genes, while here we provide evidence of alternation between multiple proliferation (IL7-R, MKI67, PCNA, PTMA, Figure 2a) and recombination genes (RAG1/2, DNTT, ADA, MS4A1/CD20, Figure 2c). This clear pattern of activity through the stages of development reveals crucial checkpoints to optimize the viability and neutralizing potential of the cell. Areas of transition between proliferation and receptor modification align with subset clustering, implying a direct relationship between successful recombination and progression in development. This checkpoint-proliferation progression is strengthened by the burst of proliferation visible in large pre-B alongside the formation of the pre-BCR (Figure 1c). Apoptosis patterns are not as disjunct and are more pervasive throughout lymphopoiesis, suggesting a more continuous regulation of non-functional B cells (Figure 2b). Past research’s suggestion that ubiquitination and subsequent apoptosis are necessary for the production of intermediate B cells, alongside the continuous patterns of apoptosis throughout development shown here, highlight the importance of cell death at every step of development (Geng et al. 2024), Figure 2)

### Previously undercharacterized pro-B cells elucidate intricacies of heavy chain recombination signaling

While pro-B is a previously identified subset of maturing B cells, this dataset is the largest scRNA seq representation of pro-B cells from human donors. It has previously been shown that developing B cells undergo rapid proliferation following successful heavy chain recombination, triggered by pre-BCR signaling (Hess et al. 2001). We show that developing B cells also undergo an earlier proliferative burst immediately following DH-JH recombination, which to our knowledge has not been described. It is unclear what initiates proliferation at this stage, as the heavy chain has not fully recombined and pre-BCR signaling is presumably not possible. Differential expression analysis between cells proliferating after D-J vs V-DJ recombination events reveals upregulation of a variety of genes previously not directly linked to B cell development (NPY, LTB, CYGB, SOCS2, LCN6, BAALC, EGFL7, COL5A1, IFITM3, SMIM3, KLF10, LST1, SYNGR1, and JCHAIN), indicating that the regulatory networks involved in proliferation during development remain to be detangled. In exploring possible signals triggering proliferation in pro-B cells, JCHAIN specifically correlated with the region of cells transitioning between D-J recombination and proliferation, possibly implicating it as a potential signaling molecule. While this is a non-canonical context for JCHAIN, recent research has suggested an evolutionary link between it and the CXCL family of genes, which are commonly utilized chemokines in B cell development (Kawasaki et al. 2024).

### VDJ sequencing reveals per subset modification of V-J pairing frequency

The defined developmental steps by which V(D)J segment and clonotype frequency are modulated to achieve such combinatorial diversity through development remain obscure (Figure 3b). Processes such as pre-BCR signaling in the pro-B phase, receptor editing in the pre-B phase, and central tolerance in intermediate B cells all have the potential to rearrange the CDRH3 region and promote clonal deletion (Winkler and Mårtensson 2018; Korzhenevich et al. 2023; Shimizu, Sun, and Ohnishi 2024). Furthermore, the particular B cells which are encouraged to proliferate depend on various checkpoints throughout development. Here, we show how the stage of development from which a cell originates affects the probability of pairing between variable and joining segments of the CDRH3 (Figure 3a). Observed differences in CDR between subsets highlights the process of receptor optimization throughout development. Understanding the processes which control V gene frequency in particular could simplify the process of B cell and antibody development, modeling, and fabrication.

### Models of VDJ pairing highlight heavy and light BCR segment contributions to diversity and development

Predicting aspects of the BCR is particularly complex as cells are developing because developmental subsets as defined here are essentially snapshots of different cells (as opposed to the same cells captured throughout development). Interestingly, predicting individual donors from V_H_/J_H_ pairs is more feasible than predicting the subset the sequence originated from, implying that repertoires between individuals are more distinct than between developmental phases, likely due to maturation processes outside of the bone marrow (Briney et al. 2019) (Figure 4). Pairing scores of variable regions from paired heavy and light chains demonstrate the evolution of the BCR throughout development, with the large pre-B pairing score distribution showing the most unlikely pairings and the small pre-B to intermediate B cell distribution shift highlighting migration of sequences towards more similar mature BCRs (Guo et al. 2025) (Figure 4a). However, SVM subset predictions are not more reliable in older cells, with V_H_/V_L_ pairs from intermediate B cells conferring the least confident subset predictions (Figure 4b and c). The difference between subsets in both ROC-AUC and ImmunoMatch pairing analyses provide some clarity to the stepwise refinement of heavy and light chain pairing and the contribution of sequence pairs to repertoire diversity even before cells leave the bone marrow, however subset prediction seems to require more nuance than CDR3 gene usage. In combination with gene expression data, ML approaches to modeling B cell development have the potential to provide a more complete reference for the processes and checkpoints necessary for repertoire diversity.

### Depth of expression data reinforces our understanding of B cell development

While scRNA seq is commonly used to observe the process of B cell maturation, this evaluation of immune cell development from human bone marrow is unique in its size and granularity ((King et al. 2021)1, (Xiao et al. 2024), (Lee et al. 2021)). Our evaluation of the transcriptome shows the timeline of how cells navigate from pluripotency to intermediate B cells and reveals insights on genetic markers and checkpoints necessary for immunological functionality (Figure 5). Patterns of expression are consistent with past research but also show in more detail how cells proceed through commitment to B cell lineage, V(D)J recombination, and clonal expansion (Figure 1c). The ontological analysis of these data provides additional clarity to the narrative of lymphopoiesis from pluripotency to intermediate B cells (Figure 5). Enriched terms in each subset highlight the checkpoints of B cell differentiation, pre-BCR signaling, and eventual MHC class II expression, strengthening our understanding of B cell developmental phase trajectory. When examining gene expression differences before and after the formation of the pre-BCR, DEG ontology confirms preestablished signal cascades which associate the formation of the pre-BCR with increase of cell division (Figure 6a,7b) (Herzog, Reth, and Jumaa 2009). It appears as though the formation and expression of the pre-BCR is the most significant checkpoint in B cell lymphopoiesis, triggering the largest proliferation event throughout development (Figure 1c, Figure 3a, Figure 6a, 7b).

These findings highlight the ongoing modifications to the CDRH3 sequence which proceed through B cell development, namely sequence shortening and optimization of particular V-J combinations. The transcriptional landscape of development presented here shows how expression and BCR formation patterns strengthen our understanding of B lymphopoiesis and heighten our capability of modeling and thus producing functional antibodies.

## AUTHOR CONTRIBUTIONS

CD, JH and BB conceptualized the study. Data generation was performed by JH. Data analysis was performed by CD and JH. The manuscript was prepared, revised and reviewed by all authors.

## FUNDING

This work was funded by the National Institutes of Health (P01-AI177683, U19-AI135995, R01-AI171438, P30-AI036214, and UM1-AI144462) and the Pendleton Foundation.

## DECLARATION OF INTERESTS

BB is an equity shareholder in Infinimmune and a member of their Scientific Advisory Board.

